# Label-free multiplex microscopic imaging by image-to-image translation overcoming the trade-off between pixel- and image-level similarity

**DOI:** 10.1101/2024.11.25.625310

**Authors:** Takashi Morikura, Akira Funahashi

**Affiliations:** Graduate School of Fundamental Science and Technology, Keio University, Kanagawa, Japan; Department of Biosciences and Informatics, Keio University, Kanagawa, Japan

## Abstract

Establishment of multiplex microscopic imaging without labeling is indispensable for understanding complex interactions of subcellular components. Toward the establishment of label-free multiplex microscopic imaging, image-to-image translation models that extract images of multiple subcellular components from bright-field images via nonlinear processing have attracted attention. However, the performance of conventional models is limited by a trade-off relationship between pixel- and image-level similarity, which degrades imaging performance. Here, we developed an image-to-image Wasserstein Schrödinger Bridge model to achieve high-performance image-to-image translation at the pixel level using Schrödinger Bridge while minimizing Wasserstein distance at the image level. Our model dramatically outperformed the conventional models at both levels simultaneously, reducing the mean squared error by 410-fold and improving the structural similarity index measure by 17.1-fold. Label-free multiplex microscopic imaging based on our model paves a way for the analysis of the interactions of subcellular components.

## Introduction

Biological systems in cells are emergent by a spatiotemporal interaction of subcellular components [1, 2, 3], and multiplex microscopic imaging allows observation of these components simultaneously [4]. Fluorescence multiplex microscopic imaging of specifically labeled subcellular components is widely used and has contributed to the development of the biological and medical fields, including molecular biology, biophysics and drug discovery screening [5, 6, 7]. However, fluorescence imaging has disadvantages such as photobleaching and phototoxicity [8, 9], and the labeling substances may interfere with the natural dynamics of the studied object [10]. Multiplex microscopic imaging without labeling is therefore needed.

Toward the establishment of label-free multiplex microscopic imaging, deep learning-based image-to-image translation models have attracted attention [11, 12]. These models aim to extract multiple images of subcellular components from bright-field images via nonlinear image processing [13]. These models are trained to solve a regression problem with bright-field images of cells as input and fluorescence images of subcellular components as output. In image-to-image translation, it is essential to achieve a balance between structural similarity, which reflects pixel-level transformation performance, and distribution similarity, which reflects image-level transformation performance.

Conditional generative adversarial networks (cGAN) have been proposed to achieve high structural similarity at the pixel level [14]. cGAN consist of two neural networks: a generator is trained to predict output images from input images, and a discriminator is trained to discriminate the predicted images and ground truth images. By training the generator to predict high-quality images that can fool the discriminator, high-performance image-to-image translation at the pixel level can be achieved. On the other hand, due to the deterministic nature of cGAN, the generated images are of low diversity, which makes it difficult to accurately capture the image-level distribution [15].

To increase the diversity in the cGAN-based image-to-image translation model, a conditional diffusion model has been proposed that incorporates a stochastic Markov process [16]. A diffusion model that is widely used today as generative artificial intelligence learns de-noising processes so that output images can be predicted from noisy images followed by a Gaussian distribution [17, 18]. In the conditional diffusion model, the input images are used as conditions in de-noising processes to achieve image-to-image translation indirectly [16]. Due to its stochastic nature, the generated images are of high diversity, making it well-suited for capturing image-level distributions [19, 20]. On the other hand, because it cannot learn image-to-image translation directly, the pixel-level structural similarity is insufficient.

The image-to-image Schrödinger Bridge model (I^2^SB) has been proposed as an extended conditional diffusion model that can learn image-to-image translation directly [21]; it learns stochastic translation directly from the input images to the output images using the Schrödinger Bridge framework. It is expected to achieve high structural similarity at the pixel level by directly learning the translation and high distribution similarity at image-level owing to its stochastic nature. If stochastic translation is considered as a path, the training process of this model is equivalent to the problem of minimizing the Kullback–Leibler divergence between the ground truth path distribution and the predicted path distribution [22, 23]. However, because the Kullback–Leibler divergence is not a distance measure, it cannot precisely reflect the geometric structure of the data distribution [24]. In the worst cases, the divergence diverges where the ground truth and predicted distribution do not overlap and a discontinuous jump occurs near the optimal parameter [25]. These intrinsic nature of Kullback–Leibler divergence may make it difficult to improve the translation performance.

Recently, the Wasserstein distance has attracted attention as a natural distance measure that overcomes the issues of the Kullback–Leibler divergence [24]. The Wasserstein distance, an optimal transport distance, is defined as the infimum of moving costs between distributions. Fortunately, the Wasserstein distance is continuous and differentiable almost everywhere [25]. Theoretical analysis showed that the upper bound of the generalization error in the deep-learning models trained using the Wasserstein distance may be tighter than that using the Kullback–Leibler divergence [26, 27]. Models trained using the Wasserstein distance have better generative performance than models trained using Fisher metrics, including the Kullback–Leibler divergence [25], but the computation to calculate the Wasserstein distance by minimizing moving costs between high-dimensional distributions such as those that generate images is intractable [28, 29]. We focused on the conditional Wasserstein generative adversarial network with a gradient penalty (cWGAN-GP) framework [30, 31] as a solver that computes the Wasserstein distance approximately. The cWGAN-GP framework uses the Kantorovich–Rubinstein representation, which is a dual formulation of the Wasserstein distance computation. This dual formulation by using Lipschitz continuity constraints decouples the calculation of combinations between the distributions in the Wasserstein distance computation, resulting in reducing the computational costs. We hypothesized that if we could stabilize the training of the I^2^SB model by replacing the minimization problems of the Kullback–Leibler divergence with minimization problems of the approximate Wasserstein distance using the cWGAN-GP framework, we would be able to develop a high-performance image-to-image translation model to be used in quantitative label-free multiplex microscopic imaging.

In this study, we developed an image-to-image Wasserstein Schrödinger Bridge model. Our model was designed to achieve high-performance image-to-image translation at the pixel level while minimizing the Wasserstein distance at the image level, and thus to overcome the trade-off constraints for establishing quantitative label-free multiplex microscopic imaging. Using published datasets [32], we evaluated the performance of our model in image-to-image translation to extract images of subcellular components from *z*-stacked, bright-field images. Our model dramatically outperformed the conventional models in structural similarity and distribution similarity. Although our model was trained without meta-information (experimental conditions and biological information), it was robust for the experimental conditions and maintained the biological information with high-quality, which are indispensable functions in the microscopic imaging. Label-free multiplex microscopic imaging using our model provides a quantitative approach to the analysis of complex interactions in cellular biological systems.

## Results

### Pixel-level structural similarity

Our model was trained to learn the task of paired image-to-image translation by supervised learning. We used microscopy images stored in the large public dataset cpg0000-jump-pilot [32] to train the model for performing image-to-image translation. The input images were bright-field images captured in three slices along the *z*-axis, and the output images were fluorescence microscopy images of subcellular components recorded in five channels (Figure 1a, Supplementary Figure S1). The dataset was constructed using the images of the human osteosarcoma cell line U2OS at 48 h of culture. To test the model, a portion of the dataset was extracted in advance as a test dataset; it consisted of biologically independent samples (Supplementary Figure S2a). The remaining dataset (excluding the test dataset) was divided into the training and validation datasets by 5-fold cross validation. The best model trained on the training dataset was selected by cross validation, and the performance of this model was evaluated using the test dataset. The illumination correction functions provided in the public dataset cpg0000-jump-pilot [32] were used to suppress uneven background illumination in the microscopy images (Supplementary Figure S2b). Detailed information about the dataset is provided in the Methods section.

**Figure 1:**
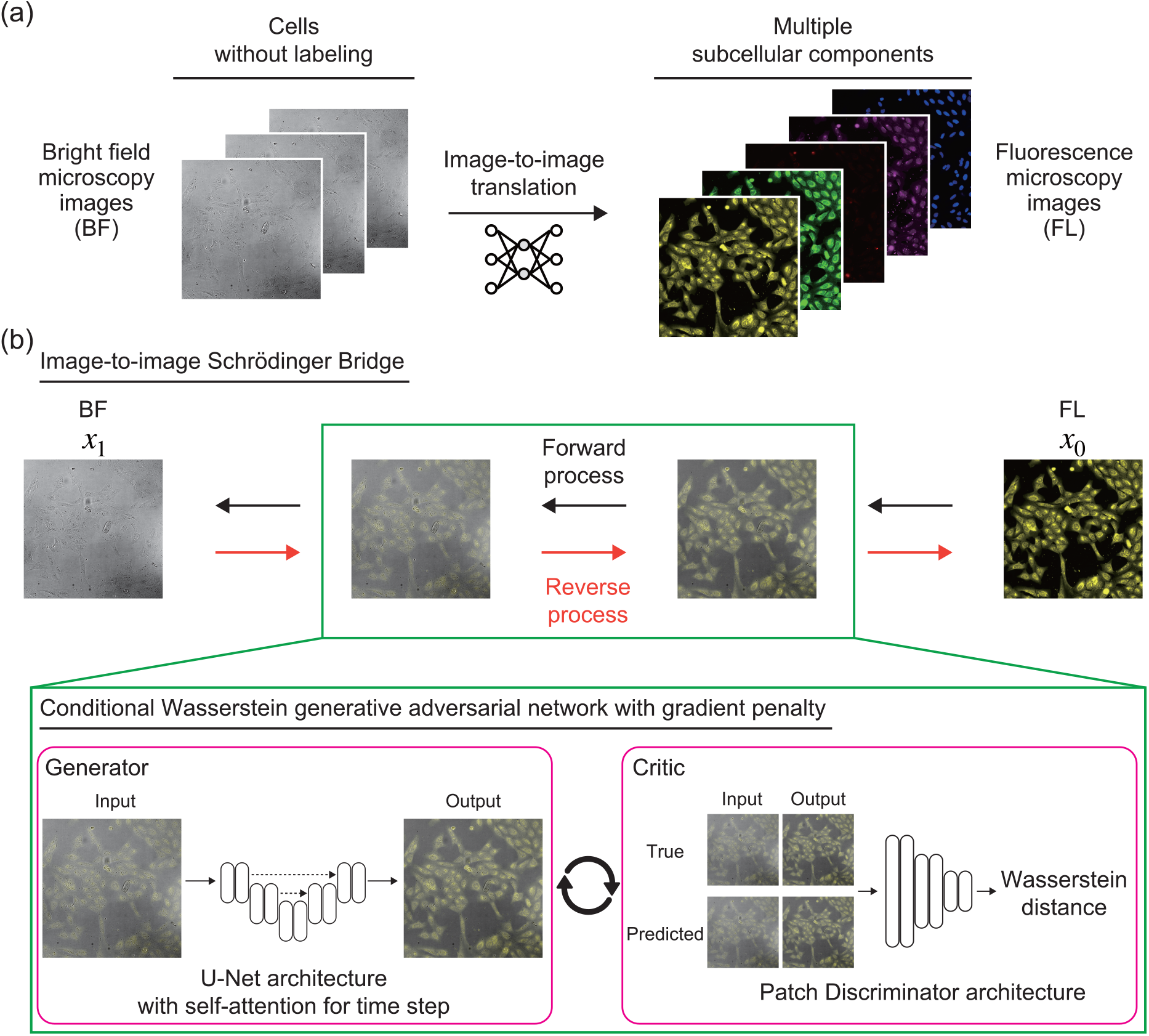
Conceptual diagram of this study. (a)Image-to-image translation by our model. The model takes bright-field images of three *z*-stacked slices as input and outputs fluorescence images of subcellular components in five channels. (b) Architecture of our model. Based on the image-to-image Schrödinger Bridge (I^2^SB) framework, the generator model learns the reverse diffusion process at each time step. During the training phase, the generator model collaborates with the critic model in the conditional Wasserstein generative adversarial network with the gradient penalty (cWGAN-GP) framework, learning to minimize the Wasserstein distance between ground truth and predicted images. The generator model is designed with a U-Net architecture with self-attention mechanisms that dynamically incorporate time-step information. The critic model follows a Patch Discriminator architecture. During the inference phase, the generator model sequentially predicts reverse diffusion to output the final predicted images. For adequate visualization, the image contrast was manually adjusted, and the images were colored using ImageJ 1.53t in this figure, whereas the images used to train our model were automatically preprocessed as described in the Methods section.

Our model consisted of the generator model of I^2^SB [21] and the critic model of cWGAN-GP [30, 31] (Figure 1b). The generator model aimed to transform images, and the critic model aimed to calculate the approximate Wasserstein distance between the generated images and ground truth images. As quantitative performance evaluation metrics of the models, we used the mean absolute error (MAE) and mean squared error (MSE), which evaluate the similarity of pixel values, structural similarity index measure (SSIM), which evaluates structural similarity by considering the spatial relationship between pixels, and peak signal-to-noise ratio (PSNR), which evaluates structural similarity on the basis of human perceptual evaluation scales. To compare the performance of our model to those of conventional models, the following three models were trained and evaluated using the same dataset: (i) Palette [33], a de facto standard conditional diffusion model in paired image-to-image translation; (ii)guided-I2I [16], a modified Palette model that takes advantage of the characteristics of microscopy images; and (iii) I^2^SB [21], a Schrödinger Bridge model (an extended conditional diffusion model). Detailed information on the architecture of our model, training conditions, and performance evaluation methods is provided in the Methods section. Palette and guided-I2I were unable to maintain cell shapes (Figure 2), whereas I^2^SB and our model maintained them with high quality. I^2^SB predicted images with low contrast, whereas our model predicted images with high contrast, similar to that in the ground truth images.

**Figure 2:**
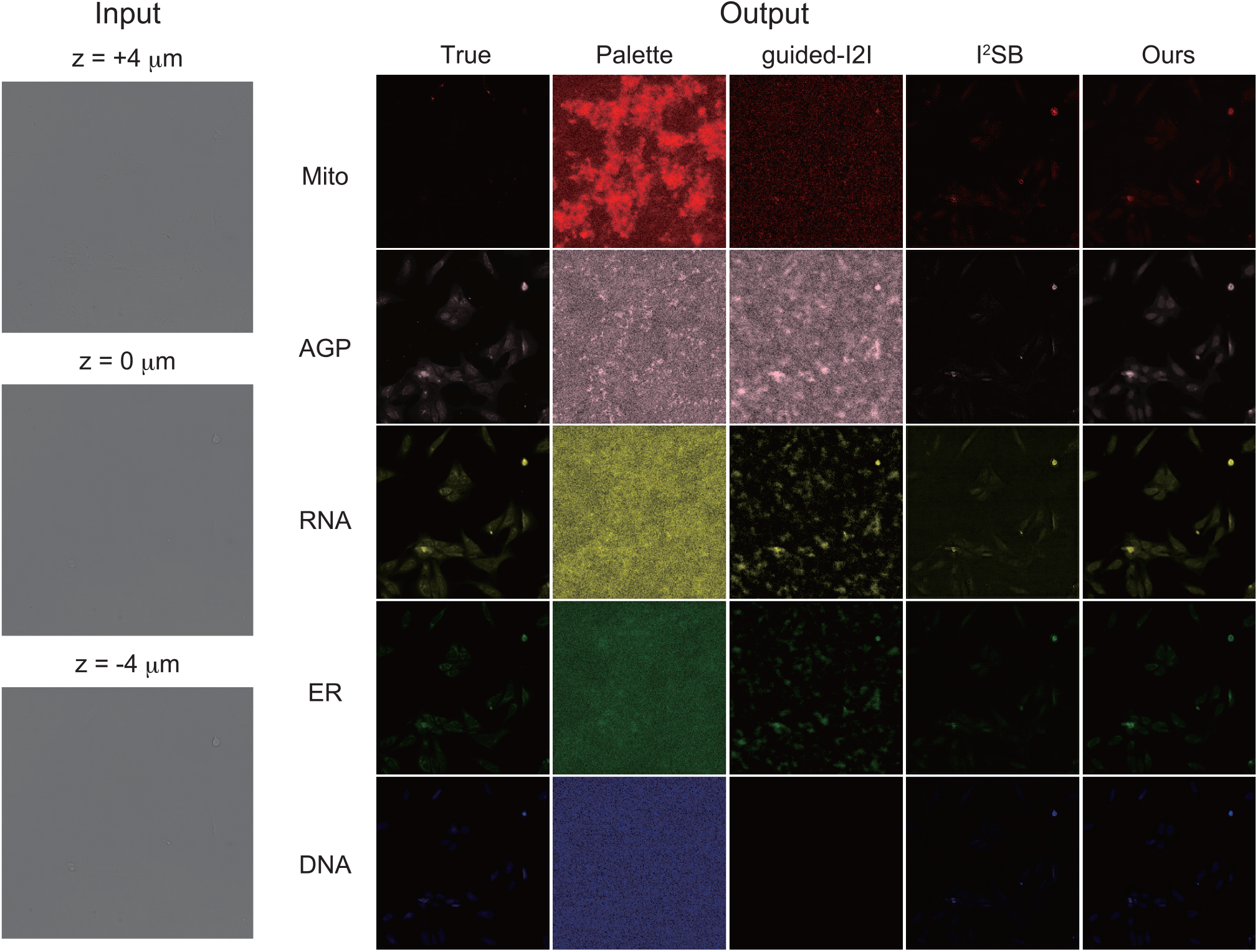
Representative images predicted by our model and conventional models. Input bright-field images were captured as three slices spaced at ±4 *µ*m along the *z*-axis. Right panel shows the output images of different models: the ground truth, Palette, guided-I2I, I^2^SB, and our model. Subcellular components (names of channels that captured them): mitochondria (Mito); Golgi, plasma membrane, and actin cytoskeleton (AGP); nucleoli and cytoplasmic RNA (RNA); endoplasmic reticulum (ER); and nucleus (DNA). The models and channel names are described in detail in the Methods section.

Quantitative evaluation showed that our model dramatically outperformed the conventional models in all evaluation metrics (Table 1). In SSIM and PSNR, which indicate pixel-level structural similarity, I^2^SB and ours dramatically outperformed the conditional diffusion models (Palette and guided-I2I). In addition, the performance of our model was significantly better than that of I^2^SB. Unlike conventional models, our model was able to maintain pixel-level structural information in images with high quality. It is worth noting that our model improved SSIM by up to 17.1-fold and PSNR by up to 3.87-fold compared to conventional models. In addition, our model also improved SSIM by 15.46 points compared to I^2^SB, which showed the best performance among conventional models, demonstrating the superiority of our model. MAE and MSE indicate pixel-level similarity; MSE of I^2^SB and ours was lower by two orders of magnitude than those of the conditional diffusion models, while MAE was of a similar order. In addition, it is worth noting that our model reduced MSE by up to 410-fold and MAE by up to 2.44-fold compared to conventional models. Our model also reduced MSE by 2.47-fold compared to I^2^SB. Thus, our model was able to produce high-quality predictions. Overall, our model dramatically improved the performance of image-to-image translation while maintaining high-quality structural similarity. Our model was able to accurately extract images of subcellular components from bright-field images without any labeling.

**Table 1:** Pixel-level structural similarity. Models were trained with images of U2OS cells at 48 h and evaluated with a test dataset (images of U2OS cells at 48 h). Columns represent the evaluation metrics. The mean values across the test dataset (n = 384) are shown. The best values are in bold. Evaluation metrics are described in the Methods section.

| Model | MAE | MSE | SSIM | PSNR |
| --- | --- | --- | --- | --- |
| Palette | $3.891 \times 10^4$ | $8.777 \times 10^8$ | 0.0487 | 8.927 |
| guided-I2I | $2.692 \times 10^4$ | $9.636 \times 10^8$ | 0.0963 | 13.31 |
| I<sup>2</sup>SB | $3.010 \times 10^4$ | $5.786 \times 10^6$ | 0.6818 | 30.12 |
| Ours | **$1.593 \times 10^4$** | **$2.347 \times 10^6$** | **0.8364** | **34.52** |

### Image-level distribution similarity

To evaluate the image-level distribution similarity, we performed dimension reduction on the ground truth images and the images predicted by each model, visualizing the data distribution that these images followed. For this purpose, we selected a linear algorithm, principal component analysis (PCA) [34], and a nonlinear algorithm, t-distributed stochastic neighbor embedding (t-SNE) [35]. Our model obtained image-level distribution accurately with both algorithms, whereas the image-level distribution obtained by conventional models differed significantly from the ground truth (PCA, Supplementary Figure S3; t-SNE, Figure 3). Importantly, I^2^SB was able to make accurate predictions at the pixel level(Table 1), but the image-level distribution captured by I^2^SB differed greatly from ground truth. On the other hand, the image-level distribution captured by our model was similar to ground truth. Our model was able to capture the image-level distribution with high quality.

**Figure 3:**
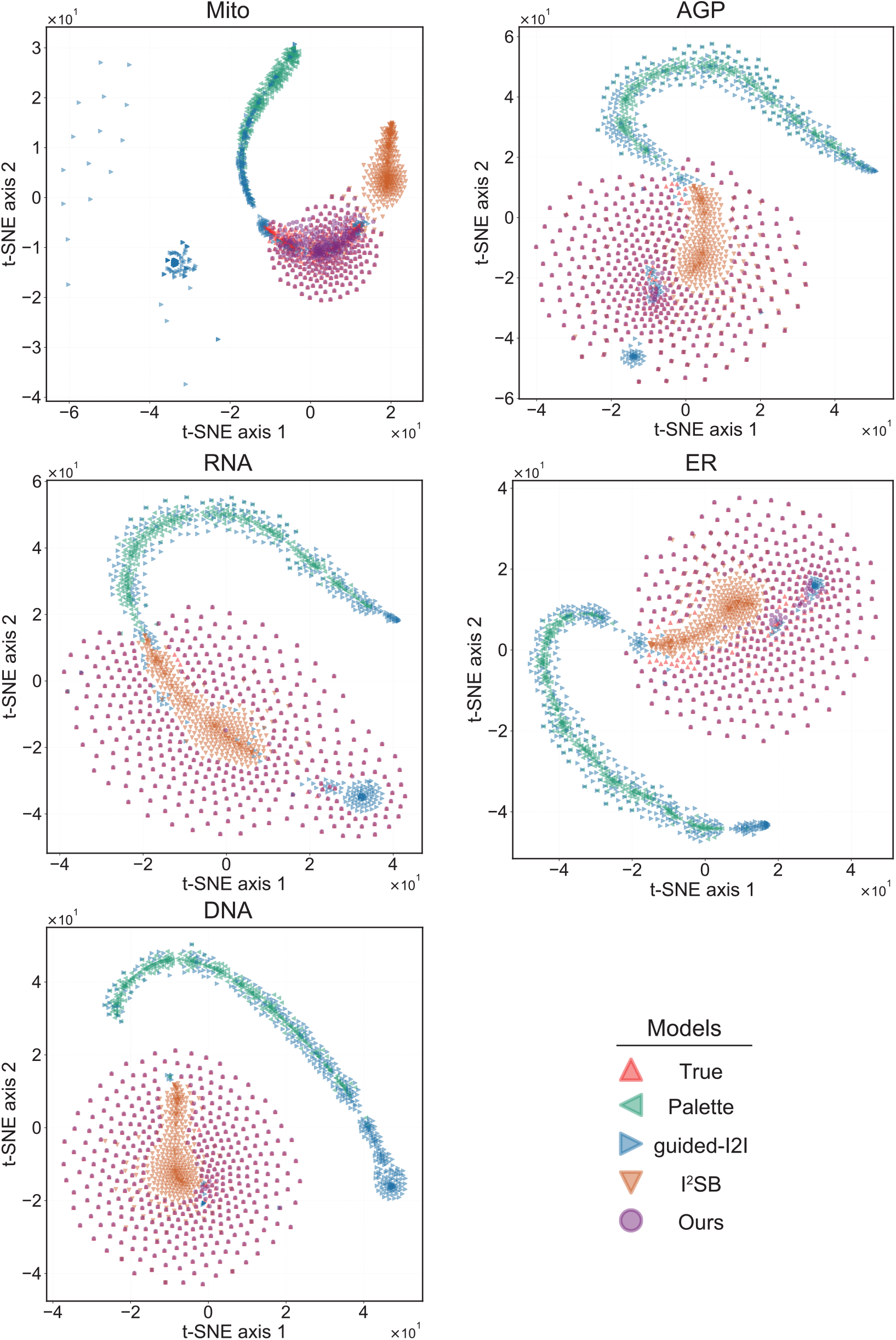
Image-level distribution similarity. Scatter plots of the feature values of the images captured in five channels and reduced to two dimensions using t-SNE.

### Robustness under different experimental conditions

To use our model as label-free imaging, the model performance must be robust under different experimental conditions. We evaluated it on images of U2OS and the human lung cell line A549 at 24 h and 48 h (Supplementary Figure S1). The models used for the evaluation were trained on images of U2OS at 48 h (Table 1). Cell density was lower at 24 h than at 48 h, and the diversity of cell morphology was higher, resulting in different spatial patterns of the subcellular components within the same cell type. Therefore, the robustness against these patterns can be evaluated by comparing model performance on images at different culture times. Since U2OS and A549 cells have different morphology and metabolic pathways, the intensity patterns differ at the same culture times, and robustness against these patterns can be evaluated by comparing model performance on images of different cell types. The model performance was evaluated for the images of U2OS cells at 24 h (Figure 4a; Extended Data Table 1), A549 cells at 48 h (Figure 4b; Extended Data Table 2), and A549 cells at 24 h (Figure 4c; Extended Data Table 3). Our model outperformed the conventional models in all evaluation metrics in all three settings, suggesting that it can robustly capture the spatial patterns and intensity patterns of the subcellular components under different experimental conditions.

**Figure 4:**
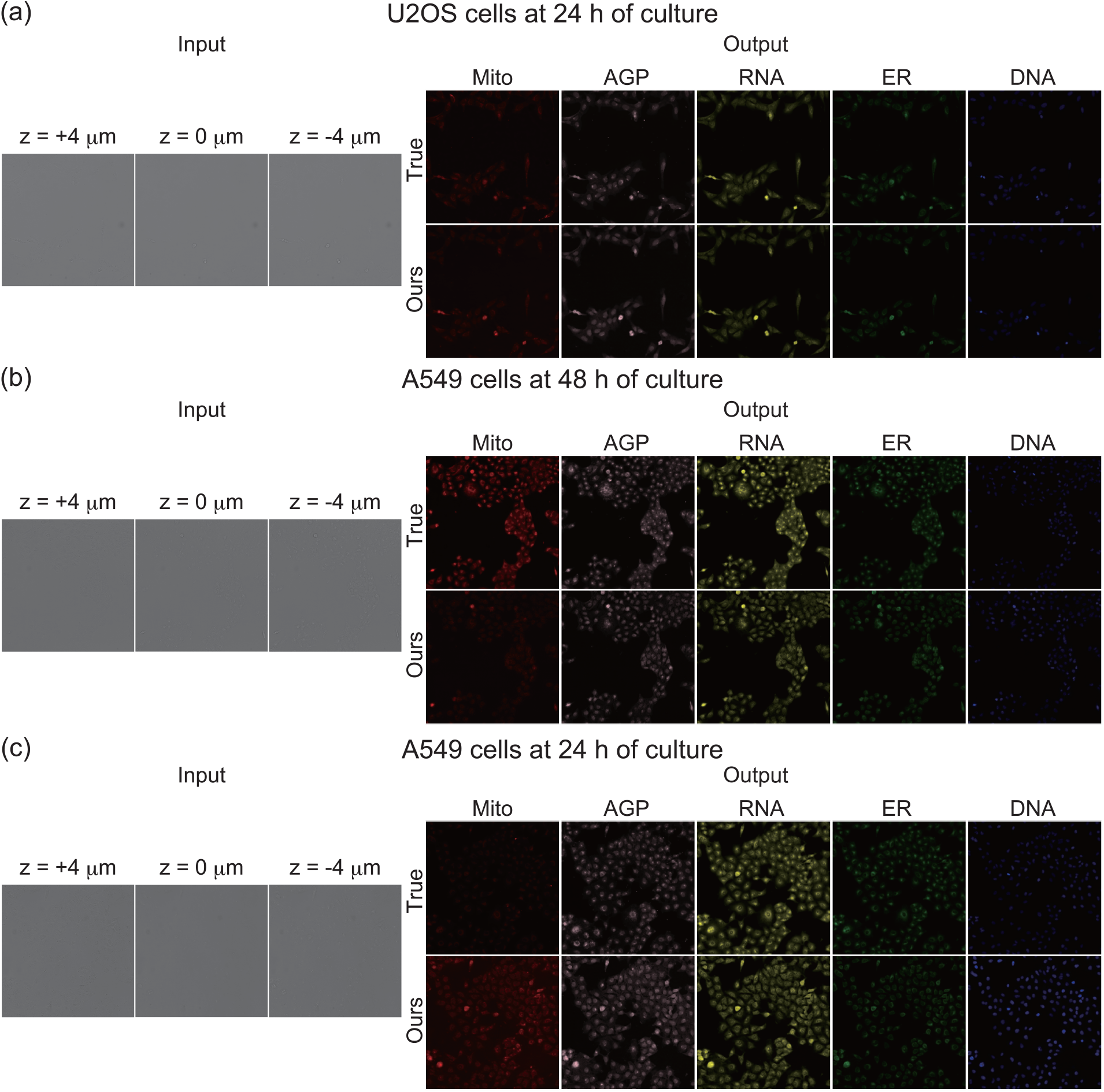
Representative images predicted by our model under different experimental conditions. The model was trained with images of U2OS cells at 48 h. Input bright-field images were captured as three slices along the *z*-axis, spaced ±4*µ*m apart. Output images include ground truth images (True) and images predicted by our model (Ours) in five channels. (a–c) Images predicted from the bright-field images of (a) U2OS cells at 24 h, A549 cells at 48 h, and (c) A549 cells at 24 h.

### Quality of biological information

To be useful for quantitative multiplex microscopic imaging, the model must precisely capture quantitative biological information. To quantify the biological information in the predicted images, we analyzed them and ground truth images in each channel using the same algorithm and compared the data between the predicted and ground truth images. We quantified information on the number of subcellular components, their shape, co-localization, and spatial patterns. Detailed information is provided in the Methods section.

To analyze the number of subcellular components, we quantitatively counted subcellular components (Figure 5a). Their number in images predicted by our model was strongly correlated with that in the ground truth images in most channels (Figure 5b); our model outperformed the conventional models (Extended Data Table 4).

**Figure 5:**
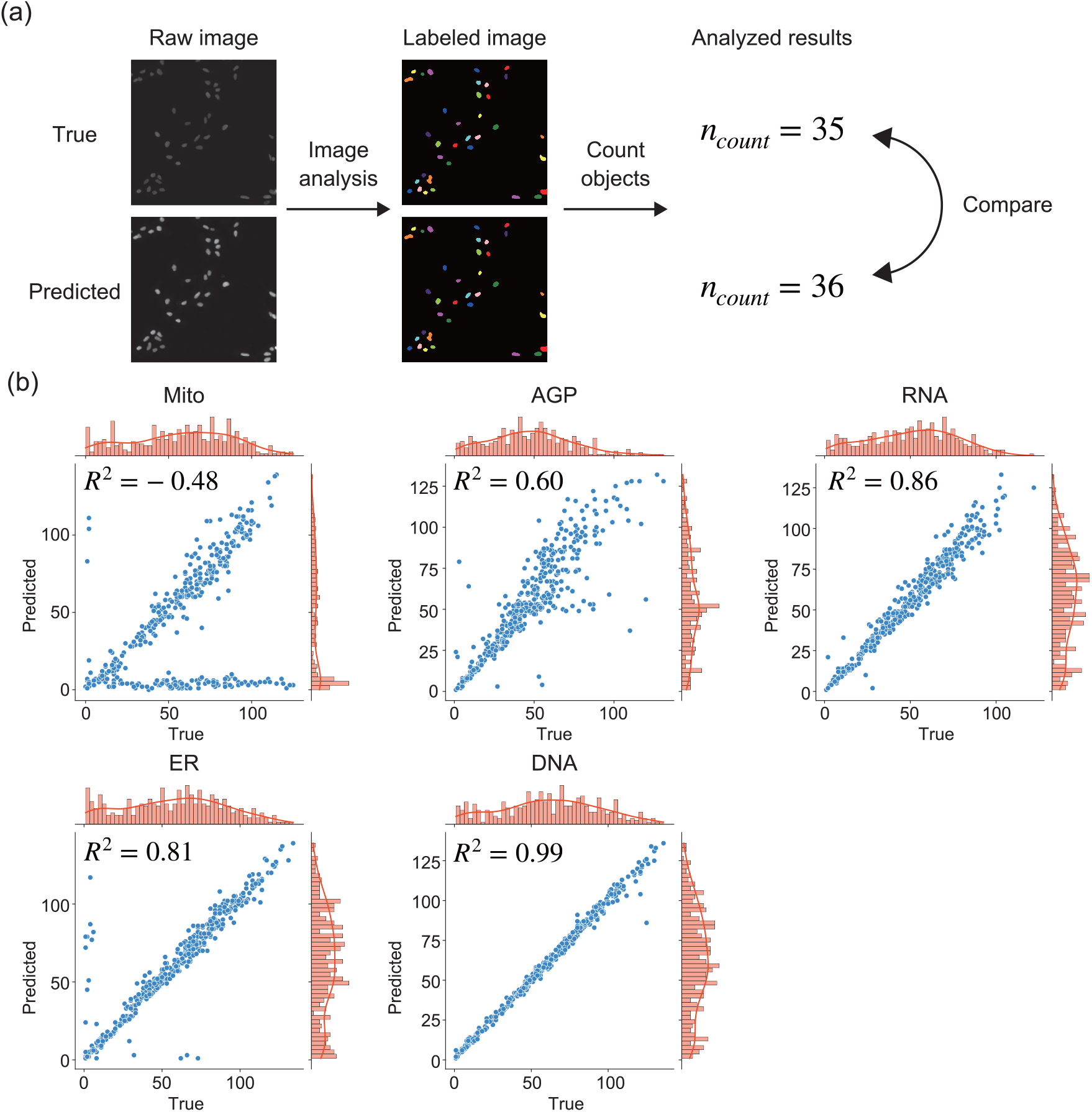
Count analysis of subcellular components. (a) Conceptual diagram. Ground truth images and predicted images were processed similarly, resulting in segmented data. Segmented objects were counted. Image processing and comparisons are described in detail in the Methods section. (b) Comparison of the counts from the ground truth images (True) and images predicted by using our model (Predicted). Channel names are shown above the plots. The right vertical axis and the upper horizontal axis show histograms (bar graphs) and kernel density (solid lines) of the counts from the ground truth and predicted images, respectively. *R*^2^, the coefficient of determination for each pair of plots.

To analyze the shape of subcellular components, we quantified the shape of subcellular components. To compare the shape of segmented objects, we applied segmentation analysis to predicted and ground truth images [36]. As quantitative shape evaluation metrics, we used IoU [37], which evaluates segmentation accuracy; SEG [38], which evaluates the absence of false-negative objects of segmentation; and MUCov [39], which evaluates the absence of false-positive objects of segmentation. In comparison with the conventional models, the shape in the images predicted by our model was more similar to that in the ground truth images (Extended Data Table 5).

To analyze co-localization of subcellular components, we calculated Manders’ colocalization coefficient (MCC) [40], which indicates the ratio of the total pixel values of a specific channel within the region of interest (ROI) to those of a different channel (Extended Data Figure 1a). MCC values for the images predicted by our model were strongly correlated with the ground truth images in most channels (Extended Data Figure 1b). In comparison with the conventional models, the MCC values for the images predicted by our model were more similar to those for the ground truth images (Extended Data Table 6).

To analyze the spatial patterns, we analyzed the intensity patterns in the spatial frequency domain of the predicted images and ground truth images. Qualitative comparison of the power spectrum in this domain showed that our model accurately captured the low-frequency components, which contain spatial structural information, rather than the high-frequency components, which contain noise (Extended Data Figure 2). The power spectrum of the images predicted by our model was most similar to that of the ground truth images (Extended Data Table 7). Although our model was trained without biological information, it captured biological information with high quality, which is indispensable in multiplex microscopic imaging. Therefore, our model has a potential to be used in multiplex microscopic imaging.

## Discussions

Label-free multiplex microscopic imaging enables simultaneous observations of different subcellular components. Recently, deep learning–based image-to-image translation models have attracted attention [11, 12], but a trade-off relationship between pixel-level structural similarity and image-level distribution similarity constrains the performance of the conventional models [15, 16]. To overcome these constraints, we developed the image-to-image Wasserstein Schrödinger Bridge model (Figure 1). The Wasserstein distance offers a natural optimal transport plan between the distributions, but it does not provide a stepwise transformation process between the images generated from each distribution. By explicitly incorporating the Wasserstein distance minimization problem into the I^2^SB framework, our model represents a stepwise transformation process explicitly from the input image to the output image by using the optimized path distribution based on Wasserstein distance learning. Using the large public dataset cpg0000-jump-pilot [32], we trained our model to extract microscopic images of the multiple subcellular components from bright-field images (Figure 2). Our model dramatically outperformed the conventional models in pixel-level structural similarity (Table 1) and in image-level distribution similarity (Figure 3, Supplementary Figure S3). Our model was robust to different experimental conditions (Figure 4, Extended Data Table 1-3) and maintained diverse biological information in different channels with high quality (Figure 5, Extended Data Figure 1-2, Extended Data Table 4-7).

Our model dramatically outperformed the conventional models in all metrics that indicate pixel-level structural similarity (Table 1). Since the test dataset used for performance evaluation consisted of samples that were biologically independent from those in the training and validation datasets, the results showed that our model can perform image-to-image translation robustly for different biological replicates. The images predicted by our model had high contrast and high quality, whereas the ones predicted by the conventional models had low contrast and low quality (Figure 2). On the other hand, in our model, false positives were observed in the Mito channel, and the images in this channel were similar to those in other channels. The loss function in our model aimed to minimize the channel-averaged approximate Wasserstein distance, which might explain the occurrence of false positives in the Mito channel unlike in the other channels. Further performance improvements are expected by modifying the loss function based on channel-specific characteristics such as the focal loss function [41].

Our model was able to obtain the image-level distribution more accurately than the conventional models (Figure 3, Supplementary Figure S3). The distribution obtained by Palette and guided-I2I differed significantly from the ground truth distribution. Although I^2^SB showed relatively high performance in terms of pixel-level structural similarity, the image-level distribution differed from the ground truth distribution. The low performance of I^2^SB as for image-level distribution the could be overcome by introducing Wasserstein distance learning. Overall, our model overcame the trade-off between pixel-level structural similarity and image-level distribution similarity and achieved high-performance image-to-image translation.

In general, the performance of deep learning–based models is considerably degraded by differences in experimental conditions under which biological samples are obtained [42]. However, our model was more robust to the perturbations caused by various experimental conditions than conventional models (Figure 4, Extended Data Table 1-3). Model robustness in the out-of-distribution is related to the diversity of model outputs [43, 44]. Considering that our model obtained high diversity, similar to that of the ground truth (Figure 3, Supplementary Figure S3), its design aimed at improving diversity might have contributed to its robustness. Because the model was not designed explicitly to improve robustness, incorporating self-learning or adversarial learning might be effective for further improvements of its robustness [45, 46, 47].

Our model captured biological information better than conventional models (Figure 5, Extended Data Figure 1-2, Extended Data Table 4-7), even though it was not explicitly given any biological information during training. Although our model captured high-quality biological information in most channels, it tended to underestimate the number and co-localization of subcellular components when it failed to predict high-quality images for some channels (Figure 5, Extended Data Figure 1). In the shape of subcellular components, our model produced false negatives, as indicated by a higher MUCov value than SEG value (Extended Data Table 5). In the spatial pattern, our model predicted incorrectly the spatial high-frequency domain representing noise (Extended Data Figure 2); it also produced false positives in some channels (Figure 2). Overall, our model tended to produce false positives in the spatial high-frequency domain, including noise, resulting in an underestimation in the process of segmentation, which is sensitive to coupling between objects. Introducing a loss function that considers the spatial frequency domain to suppress the prediction errors in the high-frequency domain [48] might improve the performance of our model.

Although we focused on a subset of biological information (Figure 5, Extended Data Figure 1-2, Extended Data Table 4-7), the fact that various analyses can be performed to extract biological information from the images predicted by our model indicates the high scalability of cell imaging using this model. For example, because conventional deep learning-based segmentation models were trained only for identifying objects in images discretely [49, 50], the conventional models cannot perform any analysis other than segmentation. On the other hand, our model was able to perform not only segmentation (Extended Data Table 5) and counting (Figure 5, Extended Data Table 4), but also co-localization analysis (Extended Data Figure 1, Extended Data Table 6) and spatial analysis (Extended Data Figure 2, Extended Data Table 7) that segmentation models cannot perform. Because the output images of our model are stored as 16-bit unsigned integer types, same as standard microscopy images, all of the analysis algorithms developed for biological microscopy images can be applied in theory. In addition, our model enables multiplex imaging without fluorescent labeling and is unaffected by photobleaching or phototoxicity, which are serious issues in conventional fluorescence microscopy [8, 9]. This feature allows an unparalleled long-term or high-speed cell imaging for the complex dynamics of multiple subcellular components, including interactions. Our model may become an essential foundational microscopic imaging technology for analyzing the spatiotemporal developmental patterns of biological multi-component and multi-body dynamics.

Our model can obtain not only high-quality semantic information such as pixel-level structural similarity and image-level distribution similarity during image-to-image transformation, but also high-quality biological information; we expect it to become a powerful tool for quantitative microscopic imaging. The use of our model in label-free multiplex microscopic imaging focused on subcellular components may contribute to a deeper understanding of complex cellular phenomena.

## Methods

### Dataset

The large public dataset cpg0000-jump-pilot [32] was used; it is stored in Cell Painting Gallery [51], the world’s largest cell microscopy database. Our model was trained on the image-to-image translation task of conversion of bright-field images of cells to fluorescence images of subcellular components (Figure 1a). The input bright-field images (three slices) were captured at ±4 *µ*m along the *z*-axis. The output images were fluorescence images of subcellular components. Detailed information about the capture conditions of these images is described in [32]. This dataset includes images of the human osteosarcoma cell line U2OS and human lung cell line A549 at 24 h and 48 h (Supplementary Figure S1). The following subcellular components stained with six fluorescent dyes were imaged in five channels: mitochondria (MitoTracker; Mito), endoplasmic reticulum (concanavalin A; ER), nucleus (Hoechst; DNA), nucleoli and cytoplasmic RNA (SYTO 14; RNA), Golgi and plasma membrane (wheat germ agglutinin; AGP), and the actin cytoskeleton (phalloidin; AGP) [32]. Cells were cultured in 384-well plates, with nine imaging regions per well. In each region, a total of eight images (three bright-field images and five fluorescence images) were captured. Four independent biological replicates were cultured in different plates on different schedules (Supplementary Figure S2a). The illumination correction functions provided in cpg0000-jump-pilot were used to suppress uneven background illumination in the images (Supplementary Figure S2b).

To train and evaluate our model, images of U2OS cells at 48 h were used. One of the four replicates was excluded in advance as the test dataset. Because the diffusion model and Schrödinger Bridge model require high computational costs for inference, we constructed 384 pairs in the test dataset by extracting one imaging region randomly from each well. The remaining dataset was split into the training and validation datasets by 5-fold cross validation. The model with the lowest MSE in the validation dataset was selected as the best candidate model in each fold. The best model (with the lowest MSE) in the whole 5-fold was selected from the best candidate models. The performance of the best model was evaluated using the test dataset. The cross-validation dataset consisted of 8295 pairs of the training dataset and 2073 pairs of the validation dataset. The total number of pairs in the datasets was 10752.

### Architecture of our model

The architecture of our model consisted of the I^2^SB [21] framework, which directly learns probabilistic transformations between images, and the cWGAN-GP [30, 31] framework, which solves the minimization problem of the Wasserstein distance between distributions (Figure 1b).

In the I^2^SB framework, a stochastic differential equation evolving over time *t* ∈ [0, 1] is considered, with the output image *X*_0_ ∼ *p*_*A*_ and the input image *X*_1_ ∼ *p*_*B*_ as boundary conditions; *p*_*A*_ is the domain distribution generating *X*_0_, and *p*_*B*_ is that generating *X*_1_. The transformation direction from *X*_0_ to *X*_1_ is defined as the forward diffusion process, whereas that from *X*_1_ to *X*_0_ is defined as the reverse diffusion process. The stochastic differential equations for each process are as follows:

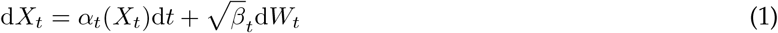

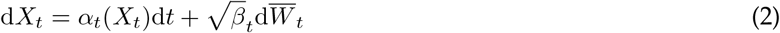

where *α*_*t*_ is the draft coefficient, *β*_*t*_ is the diffusion coefficient, *W*_*t*_ is a Wiener process, and 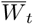 is a reverse Wiener process. In the I^2^SB framework, the Wiener process in the forward diffusion process is analytically determined, whereas the model learns the reverse Wiener process in the reverse diffusion process. When the stochastic transformation process is regarded as a path, learning proceeds in a function space of these paths to minimize the distance between the predicted path distribution and the ground true path distribution.

For the forward diffusion process, the image *X*_*t*_ at any time *t* was obtained through sampling defined as follows.

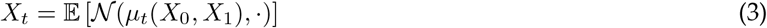

The mean *µ*_*t*_(*X*_0_, *X*_1_) of the Gaussian distribution *𝒩* (·, ·) was defined as follows (4).

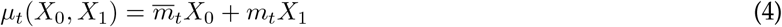

where *m*_*t*_, 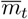 are coupling coefficients defined as 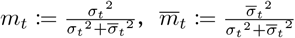, respectively. *σ*_*t*_^2^, 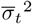 are variance defined as 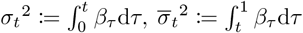 respectively.

The model learning the reverse diffusion process was referred to as a generator. As input data for the generator model, the image *X*_*t*_ at any time *t* and the input image *X*_1_ at time *t* = 1 were concatenated. On the basis of the concatenated input data, the generator model was trained to predict the ground truth noise *ϵ*_*t*_ defined as follows.

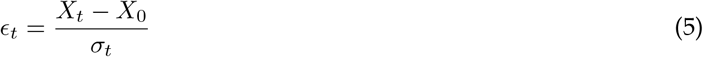

Formally, the generator was defined as 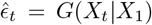. The generator consisted of a U-Net architecture with self-attention layer [21], dynamically incorporating time information. During the model training, the generator learned the reverse diffusion process at a time *t* randomly sampled from a uniform distribution *𝒰* (0, 1), enabling uniform learning of the entire reverse diffusion process over the time interval [0, 1]. In the inference phase, the generator sequentially predicts the noise 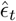 at each time *t* = *n* ∈ [0, 1], achieving the image-to-image translation from the input image *X*_1_ to the output image *X*_0_. The image-to-image translation during the inference was defined as follows.

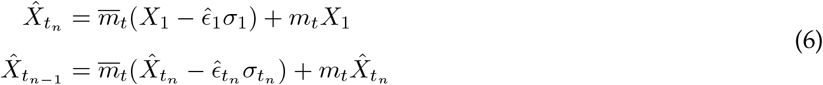

where 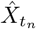 is the predicted image at time *t* = *n*, satisfying *t*_1_ = 1 *> t*_*n*_ *> t*_*n−*1_ *>* … *> t*_0_ = 0. At the time *t* = 0, the final predicted image 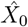 was output.

In the cWGAN-GP framework, the approximate Wasserstein distance between the distribution *p*_*r*_ of the ground truth data ***x***_*r*_, which concatenated the input and output images, and the distribution *p*_*g*_ of the generated data ***x***_*g*_ was minimized. The Wasserstein distance is continuous and differentiable almost everywhere [30]. The Wasserstein distance *W* (*p*_*r*_, *p*_*g*_) was calculated by solving the minimization problem defined in the following equation(7).

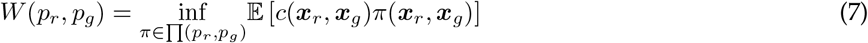

where *c*(***x***_*r*_, ***x***_*g*_) is the transportation cost from ***x***_*r*_ to ***x***_*g*_, *π*(***x***_*r*_, ***x***_*g*_) is coupling, and ∏ (*p*_*r*_, *p*_*g*_) is the set of the coupling *π* between distributions (*p*_*r*_, *p*_*g*_). The coupling *π*(***x***_*r*_, ***x***_*g*_) satisfied the following conditions: ∫*π*(***x***_*r*_, ***x***_*g*_)***x***_*g*_ = *p*_*r*_, ∫*π*(***x***_*r*_, ***x***_*g*_)***x***_*r*_ = *p*_*g*_, *π*(***x***_*r*_, ***x***_*g*_) ≥ 0. Because of the intractable computational cost to exactly compute the Wasserstein distance in the high-dimensional distributions such as images [28, 29], the cWGAN-GP framework replaced the minimization problem for computing the Wasserstein distance with a maximization problem defined as follows by using the Kantorovich–Rubinstein dual formulation [52], thereby improving the computational efficiency.

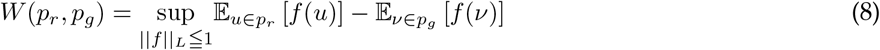

where *f* is a transport function for the ***x***_*r*_ and ***x***_*g*_, and the condition ||*f* ||_*L*_ ≦ 1 indicates that *f* is 1-Lipschitz continuous.

In our model, the Kantorovich–Rubinstein dual formulation of the Wasserstein distance was used to solve the min–max problem defined as follows.

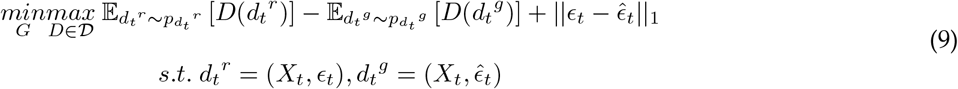

where *𝒟* is a set of function *D* that satisfies 1-Lipschitz continuity. *X*_*t*_ is the image, and *ϵ*_*t*_ is ground truth noise at time *t* in the I^2^SB framework. 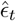 is the predicted noise defined as 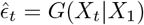. The term *d*_*t*_^*r*^ is the ground truth data concatenating *X*_*t*_ and *ϵ*_*t*_. The term *d*_*t*_^*g*^ is the predicted data concatenating *X*_*t*_ and 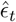. The distribution *pd*_*t*_ ^*r*^, *pd*_*t*_ ^*g*^ are the domain distributions that generate *d*_*t*_^*r*^, *d*_*t*_^*g*^. The first and second terms in equation (9) represent components for solving the maximization problem in the Kantorovich–Rubinstein dual formulation of the Wasserstein distance, while the third term represents a constraint to maintain the spatial structure of the predicted noise. In our model, the model represented by function D is referred to as the critic. The critic was trained to minimize the loss function ℒ_𝒟_ defined as follows.

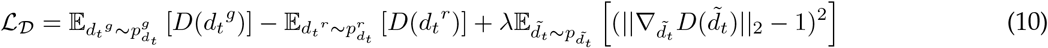

The critic was constructed with a structure similar to that of a Patch Discriminator [14]. The third term in equation(10), the gradient penalty, was a constraint to ensure that the critic satisfied the 1-Lipschitz continuity. The parameter *λ* was the weight coefficient for this constraint; its value was set to 10, as in [31]. The term 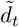 represents the interpolated data defined as follows.

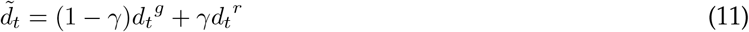

where *γ* is a random number sampled from a uniform distribution *𝒰* (0, 1). The function *G* represented the generator and the generator was trained to minimize the loss function ℒ_*𝒢*_ defined as follows.

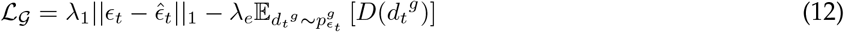

where *λ*_1_ is a weight coefficient for the L1 norm between the predicted image and the ground truth image; its value was set to 100, as in [31], *λ*_*e*_ is an adaptive weight coefficient that monotonically decreased with the training epochs. The first term represents the ℒ_1_ norm between *ϵ*_*t*_ and 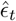. The second term represents a constraint introduced to prevent the ℒ_1_ norm of the noise from vanishing due to the adversarial loss.

In summary, our model was designed to solve the Wasserstein distance minimization problem in the I^2^SB framework by addressing the min–max problem (equation (9)).

### Training procedure

Images were stored as 16-bit unsigned integer types. At a preprocessing step, the illumination correction function provided in the datasets was applied to raw images (Supplementary Figure S2b). To reduce the computational costs, the images were resized from the original 1080 × 1080 pixels to 256 × 256 pixels. The images were normalized by min–max normalization and standardized with mean value as 0 and standard deviation value as 1, defining the range to be [−1, 1]. During training, data augmentation was performed by using random flips and random rotations. At each epoch, the input images were randomly flipped vertically or horizontally for the former and randomly rotated by 90 degrees for the latter.

Training the conditional diffusion model and Schrödinger Bridge model requires matching the dimension of the input and output data. Since the output had 5 channels, the number of channels in the input data was increased from 3 to 5. Images for two added channels were generated by mixing images captured at different positions along the *z*-axis. Since bright-field images were captured at *z* = +4 *µ*m, *z* = 0 *µ*m, and *z* = -4 *µ*m, two new channel images were generated to be closer to the images captured at *z* = +2 *µ*m and *z* = -2 *µ*m. The generation of the mixed images was defined as follows.

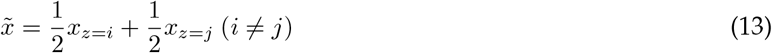

where *x* is theimage and *i, j* represent imaging positions along the *z*-axis. The generated two-channel images (mixed images captured at *z* = +4 *µ*m and *z* = 0 *µ*m or at *z* = 0 *µ*m and *z* = -4 *µ*m) were concatenated with the original three-channel input images to construct five-channel input data.

We trained the models for 100 epochs, and the mini-batch size was set to 28. The parameters in the models were optimized with Adam [53]. The hyperparameters used for training models were the same for all models (Supplementary Table S1). NVIDIA A100 80GB PCIe (FP32 19.5 TFLOPS) was used for training and inference.

### Pixel-level structural similarity

We used the MAE and MSE metrics to evaluate the pixel value similarity at each pixel; the structural similarity index measure (SSIM) [54] to evaluate spatial structural similarity; and the peak signal-to-noise ratio (PSNR) [55] to evaluate structural similarity on the basis of human perceptual quality. Each of these metrics was defined as follows.

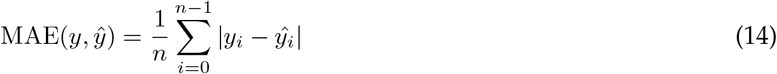

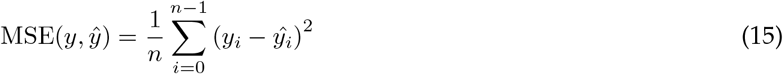

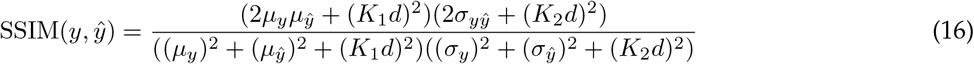

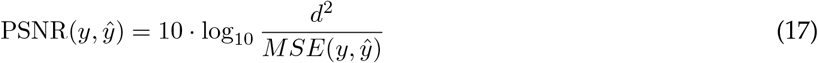

where *y* is the ground truth image; *ŷ* is the predicted image; *n* is the number of pixels; *y*_*i*_, *ŷ*_*i*_ are the pixel values at the *i*-th pixel in the ground truth and predicted image, respectively; *µ*_*y*_, *µ*_*ŷ*_ are the mean whole-pixel values in these images; *σ*_*y*_, *σ*_*ŷ*_ are the standard deviations; and *σ*_*yŷ*_ are the covariance. Constants *K*_1_ (set to 0.01) and *K*_2_ (set to 0.03) are used to avoid computational instability; *d* is the maximum pixel value in the image (set to 65535 because the images were stored as 16-bit unsigned integer type). Each metric was calculated per channel, and the mean across channels was used as the representative value for each data.

To benchmark the performance of our model, we trained and evaluated Palette [33], guided-I2I [16], and the I^2^SB model [21] on the same dataset. Palette is the *de facto* standard conditional diffusion model for paired image-to-image translation. Guided-I2I is a modification of Palette tailored to leverage the characteristics of biological microscopy images. I^2^SB is a conditional diffusion model extended by relaxing the Gaussian constraint on the input distribution. For comparative validation, the performance of the best models selected by 5-fold cross-validation was evaluated on a common test dataset.

### Image-level distribution similarity

To evaluate image-level distribution similarity, we performed the dimension reduction for the ground truth and predicted images. For this purpose, we used the linear algorithm principal component analysis (PCA) [34] and the nonlinear algorithm t-distributed stochastic neighbor embedding (t-SNE) [35]. In the latter, the perplexity, which defines the number of nearest neighbors, was fixed at 30, and the iteration number was set to 1000.

### Robustness evaluation

To evaluate the model’s robustness, we assessed its performance on images captured under different experimental conditions. The model used for evaluation was trained on images of U2OS cells at 48 h. For the evaluation, we used images of U2OS and A549 cells captured at 24 h and 48 h (Supplementary Figure S1). Differences in culture time leads to a variety of cell morphologies due to variations in cell density. Morphological differences, even within the same cell type, alter the expression patterns of subcellular components. Difference in cell type causes changes in the intensity patterns at the same culture time because of the difference of cell morphology and metabolic pathways. By using data combining different culture times and cell types, we assessed the model’s robustness to variations in expression patterns or intensity patterns of subcellular components.

### Evaluation of biological information quality in predicted images

We quantified the number, shape, co-localization, and spatial pattern of subcellular components and compared the data for the ground truth and predicted images by applying the same image processing. Each evaluation metric was calculated per channel, and the average across channels was calculated as the representative value for each data.

To count subcellular components, denoise preprocessing was performed using a median filter (kernel size: 5 pixels) and a Gaussian filter (kernel size: 5 pixels, sigma: 1.0 pixel), and then binary thresholding was performed using the Otsu algorithm [56], followed by segmentation through the watershed method to extract regions of interest. The number of the subcellular components was then quantitatively measured. Differences in the counts between the ground truth and predicted images were evaluated using MAE, MSE, and the coefficient of determination *R*^2^, which was calculated as follows.

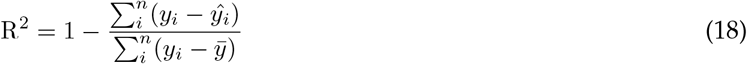

where *y* is the ground truth image, *ŷ* is the predicted image, *n* is the number of pixels, *y*_*i*_ and *ŷ*_*i*_ are the pixel value at the *i*-th pixel in the ground truth and predicted image, respectively, and 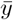 is the mean pixel value in the predicted image.

To evaluate the shape similarity of objects extracted from the ground truth and predicted images [36], we processed the images as above. To evaluate segmentation accuracy, we used the IoU metric [37]. To evaluate the absence of false-negative objects of segmentation, we used the SEG metric [38]. To evaluate the absence of false-positive objects of segmentation, we used the MUCov metric [39]. Each of these metrics is defined as follows.

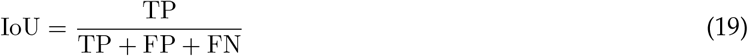

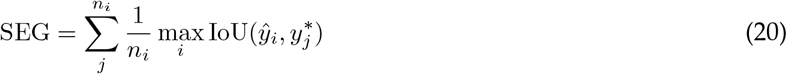

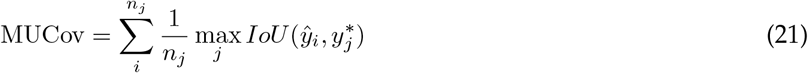

where TP is the number of pixels where the ROI in the ground truth matches that in the predicted image; FP is the number of pixels classified as ROI in the predicted image but as background in the ground truth image; FN is the number of pixels classified as ROI in the ground truth but as background in the predicted image; *n*_*i*_ and *n*_*j*_ are the numbers of ROIs in the predicted image and the ground truth image, respectively; and *y*_*i*_ and *ŷ*_*i*_ are the ROI at the index *i* in the ground truth and predicted image, respectively. SEG and MUCov were calculated only at IoU *>* 0.5 [38].

To analyze co-localization, we calculated MCC [40], which indicates the ratio of the total pixel values of a specific channel within an ROI to those of a different channel (Extended Data Figure 1a). MCC was calculated for all combinations of channels, and the average value was used as the representative value. MCC was defined as follows.

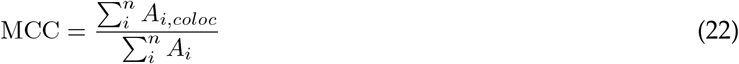

where *A*_*i*_ is the pixel value at index *i* in a specific channel, and *A*_*i,coloc*_ was defined as follows.

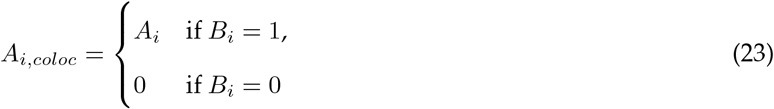

where *i* is an index number in the image; *B*_*i*_ shows whether the *i*-th pixel in the referenced channel is part of the background. If *B*_*i*_ = 1, it represents the background region, and if *B*_*i*_ = 0, it represents the ROI. To discriminate between the background and ROI, segmentation processing was done with the same algorithm used for counting. Differences in the MCC between the ground truth and predicted images were evaluated using MAE and MSE.

To examine spatial patterns, we analyzed the power spectrum (PS) in the spatial frequency domain of the ground truth and predicted images by applying the following equation to the structured data in this domain obtained through discrete Fourier transform of the images.

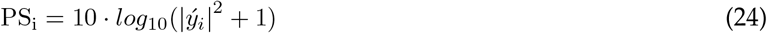

where *ý* is the structured data in the spatial frequency domain obtained by performing the discrete Fourier transform on either the ground truth or predicted image; *i* is the index in the structured data. Differences in PS between the ground truth and predicted images were evaluated using MAE and MSE.

## Code availability

The source code of this study is available from https://github.com/funalab/I2IWSB.

## Data availability

We used the dataset cpg0000-jump-pilot [32], available from the Cell Painting Gallery [51] in the Registry of Open Data on AWS (https://registry.opendata.aws/cellpainting-gallery/). The detailed structure of the datasets used for the training and evaluation in this study is available in our repository (https://github.com/funalab/I2IWSB).

## Acknowledgements

The research was funded by JST CREST, Japan Grant Number JPMJCR2331 to A.F.

## Author contributions

T.M. and A.F. designed the conceptual idea and the study. T.M. implemented the developed algorithm. T.M. and A.F. wrote the manuscript.

---

**Extended Data Table 1:** Robustness of the models to a difference in culture time. Models were trained with images of U2OS cells at 48 h and evaluated with images of U2OS cells at 24 h. Columns represent the evaluation metrics. The mean values across the test dataset (n = 384) are shown. The best values are in bold. Evaluation metrics are described in the Methods section.

| Model | MAE | MSE | SSIM | PSNR |
| --- | --- | --- | --- | --- |
| Palette | $3.930 \times 10^4$ | $8.489 \times 10^8$ | 0.0535 | 9.034 |
| guided-I2I | $2.692 \times 10^4$ | $9.590 \times 10^8$ | 0.0849 | 12.72 |
| I<sup>2</sup>SB | $3.020 \times 10^4$ | $7.649 \times 10^6$ | 0.6128 | 28.82 |
| Ours | **$1.403 \times 10^4$** | **$3.735 \times 10^6$** | **0.8048** | **32.32** |

**Extended Data Table 2:** Robustness of the models to a difference in cell type. Models were trained with images of U2OS cells at 48 h and evaluated with images of A549 cells at 48 h. Columns represent the evaluation metrics. The mean values across the test dataset (n = 384) are shown. The best values are in bold. Evaluation metrics are described in the Methods section.

| Model | MAE | MSE | SSIM | PSNR |
| --- | --- | --- | --- | --- |
| Palette | $3.931 \times 10^4$ | $8.352 \times 10^8$ | 0.0527 | 9.198 |
| guided-I2I | $2.720 \times 10^4$ | $9.087 \times 10^8$ | 0.1219 | 13.35 |
| I<sup>2</sup>SB | $3.163 \times 10^4$ | $1.586 \times 10^7$ | 0.6548 | 27.95 |
| Ours | **$2.648 \times 10^4$** | **$8.102 \times 10^6$** | **0.7698** | **30.55** |

**Extended Data Table 3:** Robustness of the models to differences in culture time and cell type. Models were trained with images of U2OS cells at 48 h and evaluated with images of A549 cells at 24 h. Columns represent the evaluation metrics. The mean values across the test dataset (n = 384) are shown. The best values are in bold. Evaluation metrics are described in the Methods section.

| Model | MAE | MSE | SSIM | PSNR |
| --- | --- | --- | --- | --- |
| Palette | $3.954 \times 10^4$ | $8.126 \times 10^8$ | 0.0647 | 9.280 |
| guided-I2I | $2.691 \times 10^4$ | $9.214 \times 10^8$ | 0.0923 | 12.40 |
| I<sup>2</sup>SB | $2.643 \times 10^4$ | $1.690 \times 10^7$ | 0.5734 | 26.40 |
| Ours | **$1.537 \times 10^4$** | **$9.136 \times 10^6$** | **0.7656** | **28.49** |

**Extended Data Table 4:**
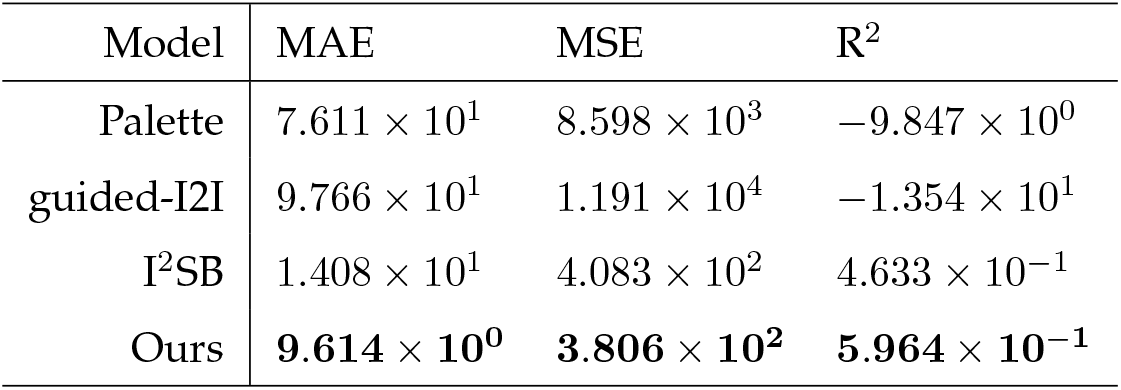
Model performance in count analysis. Subcellular components were counted in each image. Columns represent the evaluation metrics. The mean values across the test dataset (n = 384) are shown. The best values are in bold. Evaluation metrics are described in detail in the Methods section.

**Extended Data Table 5:** Model performance in shape analysis. The shape of subcellular components in each image was analyzed. Columns represent the evaluation metrics. The mean values across the test dataset (n = 384) are shown. The best values are in bold. Evaluation metrics are described in the Methods section.

| Model | IoU | SEG | MUCov |
| --- | --- | --- | --- |
| Palette | 0.2881 | 0.3566 | 0.2061 |
| guided-I2I | 0.2232 | 0.3118 | 0.1402 |
| I<sup>2</sup>SB | 0.5810 | 0.6951 | 0.5249 |
| Ours | **0.8259** | **0.7136** | **0.7336** |

**Extended Data Figure 1:**
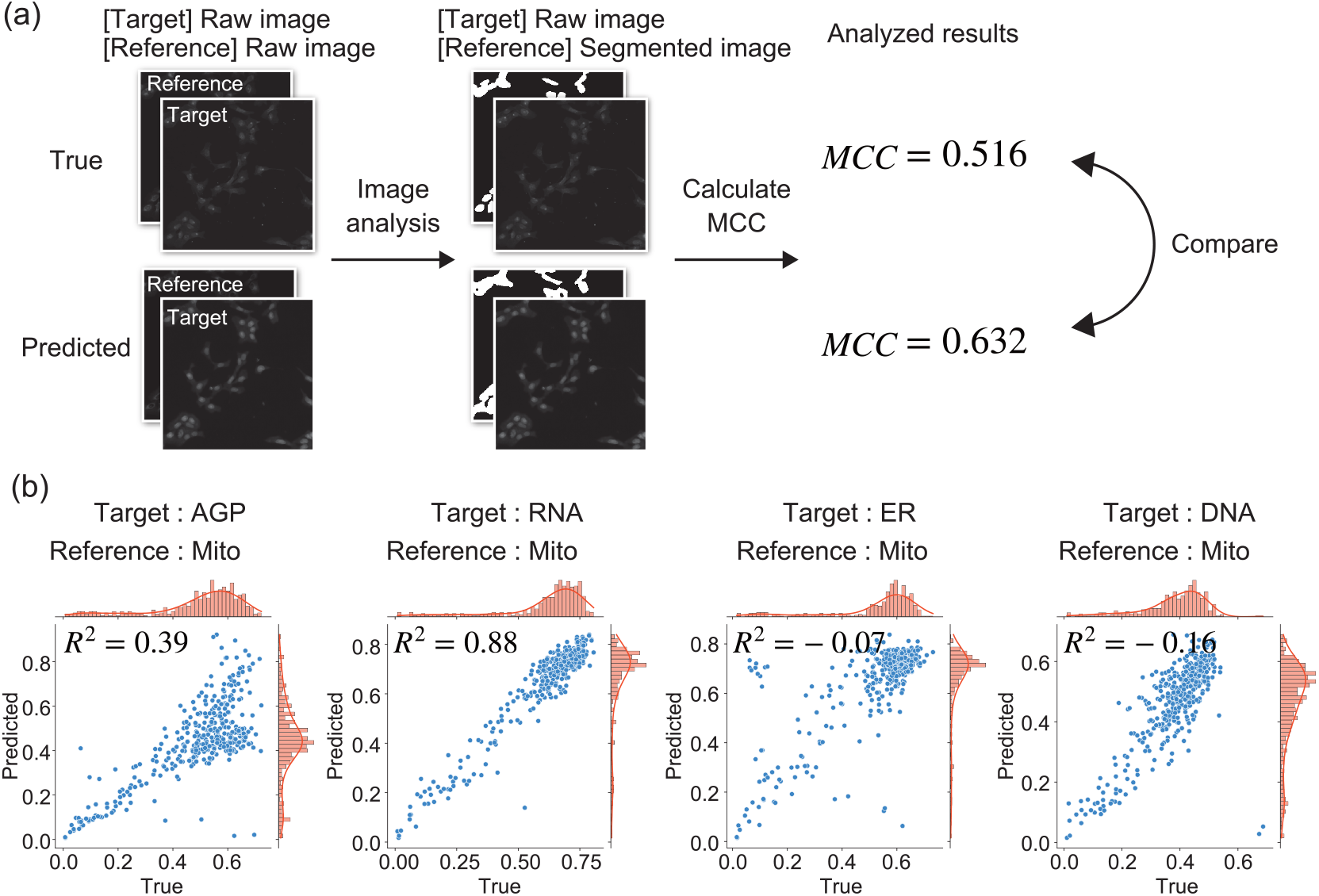
Co-localization analysis of different subcellular components. (a) Conceptual diagram. Ground truth images and predicted images were processed similarly for both the target channel and the reference channel. Manders’ colocalization coefficient (MCC) was calculated between different channels. Detailed information is provided in the Methods section. (b) Representative scatter plots showing MCC for the ground truth images and for those predicted by our model, with Mito as the reference channel. The left vertical and the lower horizontal axes represent the MCC values from the predicted and ground truth images, respectively. The right vertical axis and the upper horizontal axis show histograms (bar graphs) and kernel density (solid lines) of the MCC values from the ground truth and predicted images, respectively. *R*^2^, the coefficient of determination for each pair of plots.

**Extended Data Table 6:** Model performance in co-localization analysis. The co-localization of subcellular components was compared. MCC was calculated for all combinations of channels, and the average value was used as the representative value. Columns represent the evaluation metrics. The mean values across the test dataset (n = 384) are shown. The best values are in bold. Evaluation metrics are described in detail in the Methods section.

| Model | MAE | MSE |
| --- | --- | --- |
| Palette | 0.0508 | 0.1847 |
| guided-I2I | 0.0869 | 0.2302 |
| I<sup>2</sup>SB | 0.0850 | 0.2412 |
| Ours | **0.0201** | **0.1013** |

**Extended Data Figure 2:**
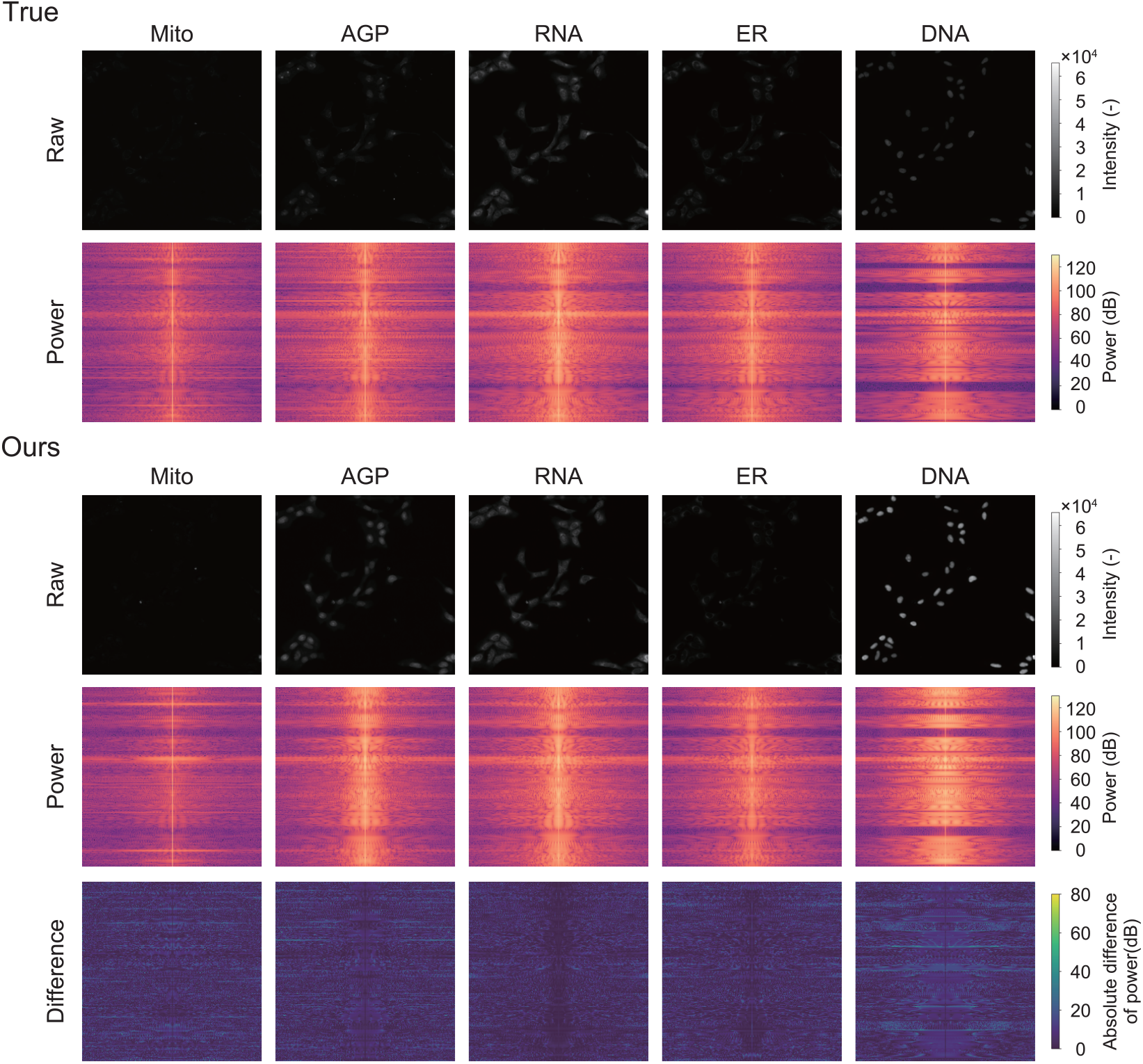
Spatial pattern analysis in the spatial frequency domain. Representative power spectra are shown for the ground truth images (True) and the images predicted by our model (Ours). The algorithm used to calculate the power spectrum is described in detail in the Methods section. The bottom section visualizes absolute differences in power spectra between the ground truth and the predicted images. The names of the channels are shown above the raw images. The central region of the power spectrum shows the low-frequency components of spatial frequency, and the outer regions show the high-frequency components.

**Extended Data Table 7:**
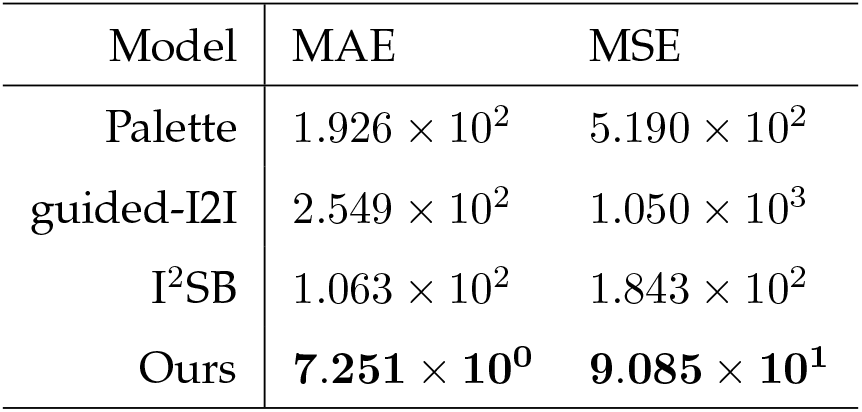
Model performance in spatial pattern analysis. The power spectrum in the spatial frequency domain of each image was compared. Columns represent the evaluation metrics. Mean values across the test dataset (n = 384) are shown. The best values are in bold. Evaluation metrics are described in detail in the Methods section.

| Model | MAE | MSE |
| --- | --- | --- |
| Palette | $1.926 \times 10^2$ | $5.190 \times 10^2$ |
| guided-I2I | $2.549 \times 10^2$ | $1.050 \times 10^3$ |
| I<sup>2</sup>SB | $1.063 \times 10^2$ | $1.843 \times 10^2$ |
| Ours | **$7.251 \times 10^0$** | **$9.085 \times 10^1$** |

