## Supplementary Material for "Label-free multiplex microscopic imaging by image-to-image translation overcoming the trade-off between pixel- and image-level similarity"

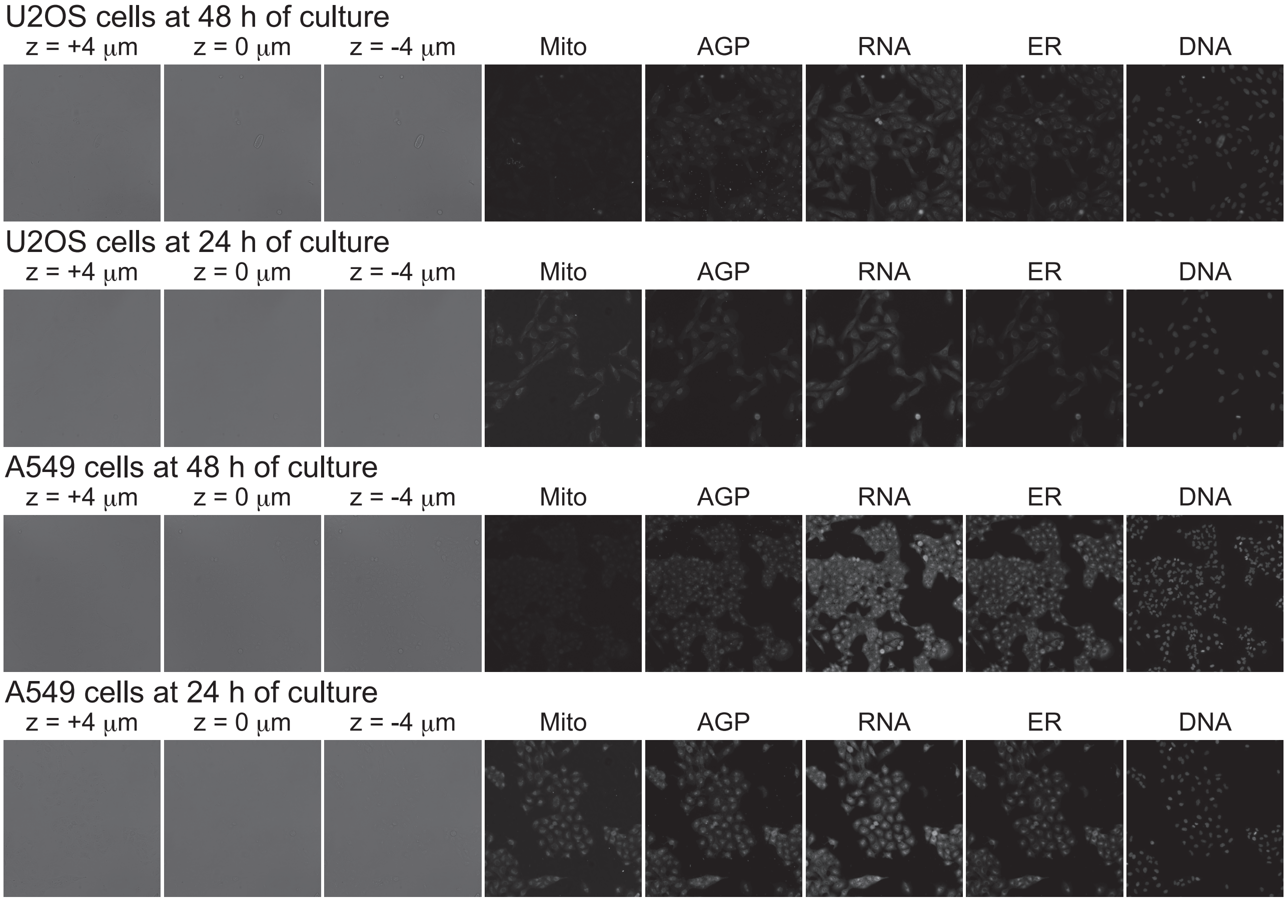


Figure S1: Representative microscopy images from the dataset.

The dataset includes images of human osteosarcoma cell line U2OS and human lung cancer cell line A549 captured at 24 h or 48 h. The input bright-field images were captured at three $z$-axis positions, spaced at $\pm 4$ $\mu$m intervals, and fluorescence images were the output. Subcellular components (corresponding fluorescence channels): mitochondria (Mito); Golgi, plasma membrane, and actin cytoskeleton (AGP); nucleoli and cytoplasmic RNA (RNA); endoplasmic reticulum (ER); and nucleus (DNA).

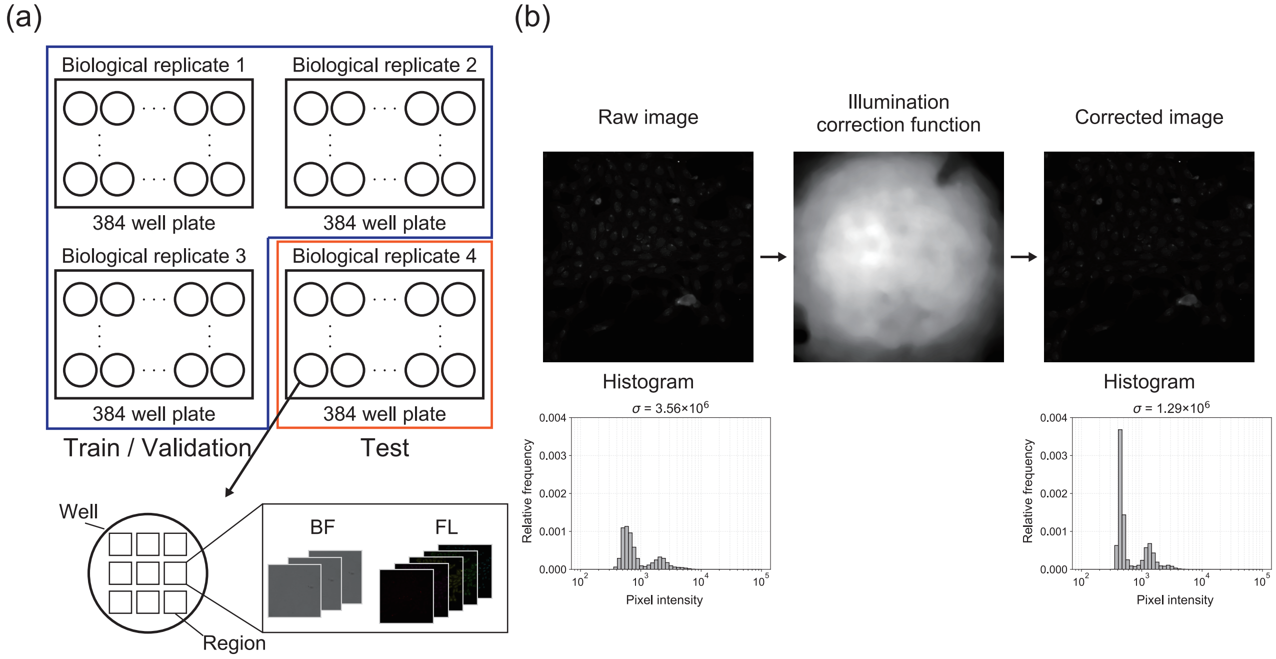


Figure S2: Conceptual diagram of the dataset and preprocessing.

(a) In this dataset, four independent biological replicates for each condition were obtained for specific cell types and culturing times. One replicate was used as the test dataset, while the remaining replicates served as the training and validation datasets. Nine imaging regions were used in each well. For each imaging region, both bright-field images (BF) and fluorescence images (FL) were captured. (b) To reduce illumination unevenness in the raw images, the illumination correction function provided with the dataset was applied. The histograms show the distribution of pixel intensity values in the raw and processed images. The pixel intensity values were transformed to a logarithmic scale. The variance of the pixel intensity values within each distribution is shown above each histogram. Illumination correction decreased variance, demonstrating a decrease in illumination unevenness across the image.

Table S1: Hyperparameters used in the training and inference for our and conventional models. The values of the hyperparameters are from previous studies. Number of epochs, the total number of training epochs; mini-batch size, the number of data samples in each mini-batch during an epoch; time interval, the number of splits in the diffusion process; learning rate, the initial learning rate used in the Adam optimizer; betas, coefficients used for calculating gradients in the Adam optimizer; weight decay, coefficients for calculating the L2 norm of the parameters; critic update frequency, the ratio at which the critic was trained in comparison with the generator in the cWGAN-GP framework.

| Model: Palette | | Model: $I^2SB$ | |
|---|---|---|---|
| Hyperparameter | Value | Hyperparameter | Value |
| Number of epochs | 100 | Number of epochs | 100 |
| Mini-batch size | 28 | Mini-batch size | 28 |
| Time interval | 2000 | Time interval | 2000 |
| Adam learning rate | $5.0 \times 10^{-5}$ | Adam learning rate | $5.0 \times 10^{-5}$ |
| Adam betas | (0.9, 0.999) | Adam betas | (0.9, 0.999) |
| Adam weight decay | 0.0 | Adam weight decay | 0.0 |
| Model: guided-I2I | | Model: Ours | |
| Hyperparameter | Value | Hyperparameter | Value |
| Number of epochs | 100 | Number of epochs | 100 |
| Mini-batch size | 28 | Mini-batch size | 28 |
| Time interval | 2000 | Time interval | 2000 |
| Adam learning rate | $8.0 \times 10^{-5}$ | Adam learning rate at critic and generator | 0.0002, $5.0 \times 10^{-5}$ |
| Adam betas | (0.9, 0.999) | Adam betas at critic and generator | (0.0, 0.9), (0.0, 0.9) |
| Adam weight decay | 0.0 | Adam weight decay at critic and generator | 0.001, 0.0 |
| | | Critic update frequency | 2 |

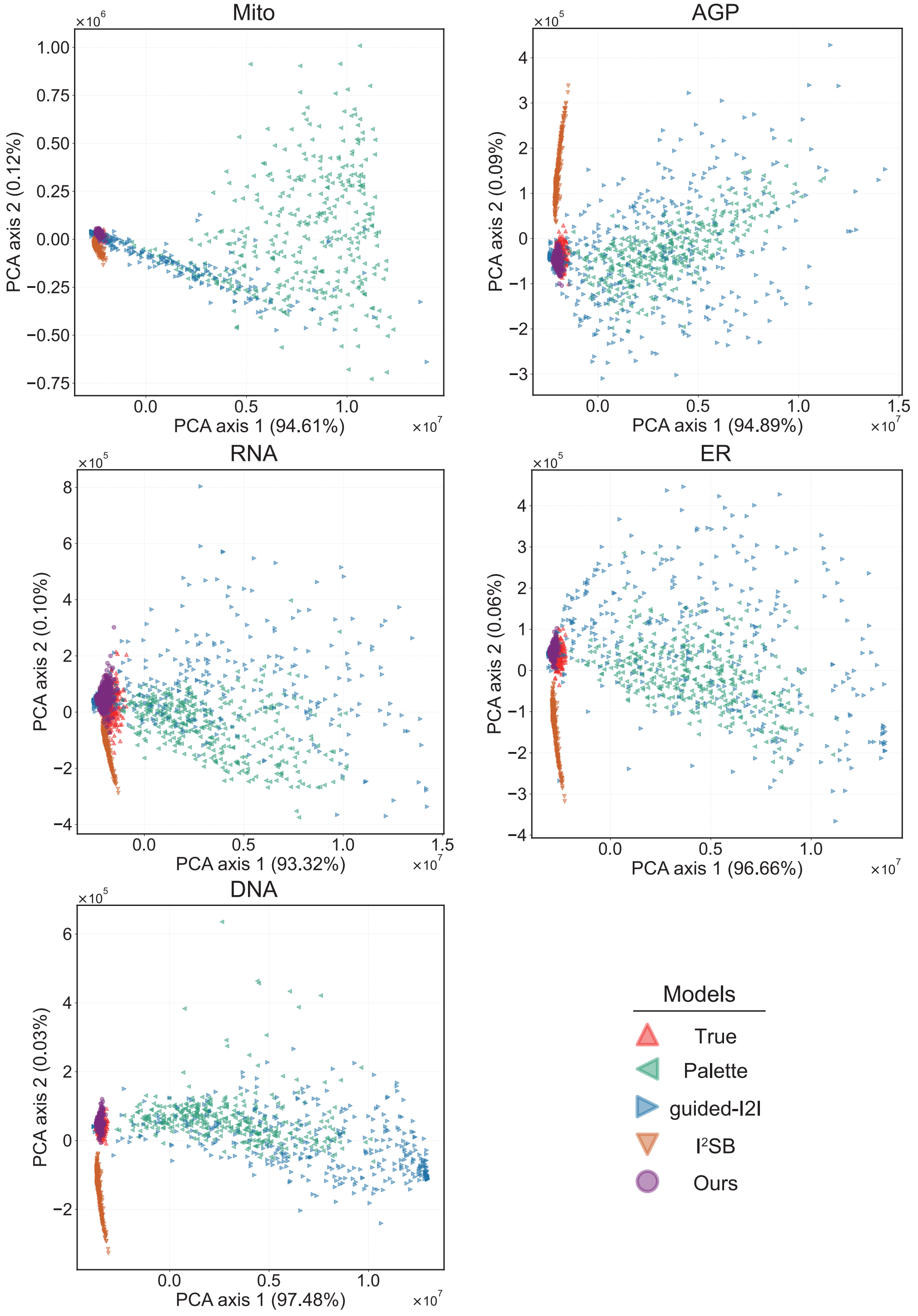


Figure S3: Scatter plots of the feature values of the images reduced to two dimensions using PCA. The labels represent each model. Channel names are above the plots.
